# Early inflammatory response mediated by Angiotensin II in cardiac arteries of normotensive mice

**DOI:** 10.1101/2023.11.06.563831

**Authors:** Thais Cristina de Souza Oliveira, Katia Aparecida da Silva Viegas, Rariane Silva de Lima, Tatiana C. Alba-Loureiro, Cintia Taniguti Lima, Luciene Cristina Gastalho Campos, Valerio Garrone Barauna, Rui Curi, Maria Claudia Irigoyen, Silvia Lacchini

## Abstract

To verify if a low dose of angiotensin II (Ang II) can induce an inflammatory response in cardiac arteries, even though blood pressure remains at normal values. Were used C57Bl/6J male mice treated with a low dose of Ang II (30ng/kg IP) and AT1R blockers. Blood pressure was recorded after 10, 30, 60min, 2 and 6hours after Saline or Ang II injection. Time curve (30 and 60min, 2, 6, 12, 24, and 48hours after treatment) for expression of inflammatory markers was evaluated in cardiac arteries (TGF-β, IL-1β, IL-6, TNF-α and ICAM-1) by immunohistochemistry and western blot. Serum TNF-α and IL-6 were analyzed by ELISA. Although Ang II did not alter blood pressure, local TGF-α and IL-6 presented an early increase in cardiac arteries. IL-1β and ICAM-1 participated in a late response to Ang II (12 hours). Ang II group showed a systemic increase of IL-6 30 to 60 minutes after treatment. AT1 and AT2 receptor blockers (losartan, 20mg/kg, and PD123.319, 15mg/kg) were given alone or concomitant to Ang II. This combination showed that Ang II increases TGF-β and IL-6 by acting on the AT1 receptor in a fast response. IL-1β regulation by Ang II seems to be mediated by both AT1 and AT2 receptors. The results suggest that, independently of hemodynamic influences, Ang II, acting on AT1R, leads to the expression of inflammatory markers in cardiac vessels. AT2 receptor seems to be involved in the counterbalance of AT1R during modulation of IL-1β production in cardiac arteries.

## Introduction

The Renin-angiotensin system (RAS) plays an important role in both physiological mechanisms of vascular control and pathological conditions. The activation of RAS after vascular injury or endothelial dysfunction has been related to vascular remodeling and atherosclerosis(Rakugi et al., 1993). Studies have shown an association between the increased expression of Angiotensin I converting enzyme (ACE) and the formation of atherosclerotic plaques (Ohishi et al., 1997, 1999). ACE is involved in major functions of the vasculature controlling the contractility, growth, and migration of vascular smooth muscle cells (VSMC), contributing to the development of hyperplasia in the tunica intima and hypertrophy in the tunica media (Dzau & Lopez-Ilasaca, 2005; Epstein et al., 1994). A previous study has demonstrated that an increase of ACE gene copies is closely related to the augment of ACE vascular activity and vascular injury in genetically modified mice (Lacchini et al., 2009). Others have shown that the levels of ACE and Angiotensin II (Ang II) are increased in the atheromatous plaque of aortic atherosclerosis and restenosis after angioplasty or stent implantation. Moreover, AT1 receptor antagonism reduces endothelial dysfunction and intimal thickening in atherosclerotic vessels (Ohishi et al., 1997, 1999; Ribichini et al., 2000). It is well described that Ang II is produced in the vasculature and its production is capable of generating intimal hyperplasia independently of hemodynamic or neurohumoral effects (Dzau & Lopez-Ilasaca, 2005; Epstein et al., 1994; Naftilan, 1994; Weir & Dzau, 1999; Ross, 1999). Considering that Ang II is associated with the proliferation and migration of VSMCs, collagen deposition, inflammation, and oxidative stress, this peptide can be described as one of the main mediators of vascular adaptation and remodeling para (Schiffrin & Touyz, 2004; Paradis & Schiffrin, 2009).

Indeed, several studies have linked the action of Ang II-AT1 receptors to the development of vascular diseases, whereas Ang II-AT2 receptors have been shown opposing effects (Navar et al., 2002, 2003). Also, it has been demonstrated that hypertensive patients exhibit increased serum adhesion molecules and inflammatory cells, which may be directly associated with the inflammatory actions of Ang II (Ruiz-Ortega et al., 2001, 2007). Besides being a hormone that plays a central role in cardiovascular homeostasis by regulating vasoconstriction and blood pressure, Ang II is considered a multifunctional cytokine with non-hemodynamic properties, modulating other cytokines as well as coagulation and growth factors (Taubman, 2003; Wolf et al., 2003). Recently, it was shown that a low dose of Ang II (which does not alter the blood pressure) induces the expression of inflammatory markers and migration of CD45 positive cells into the aorta of normotensive mice (Lima et al., 2019). Thus, the precise role and the direct activity of inflammatory cytokines in vascular diseases are still unknown.

As shown in the literature, Ang II is strongly associated with hypertension in clinical and experimental studies. However, such an association may represent a confusing factor if considering that hypertension per se is capable of inducing inflammatory mechanisms (Harwani, 2018; Jafri & Ormiston, 2017). Therefore, the possibility of evaluating Ang II-directly mediated signaling from blood pressure is important to understand each moment in the remodeling process and to analyze the transition from the healthy to a pathological condition.

The correlation between the inflammatory process and the physiological action of Ang II, as well as molecular mechanisms involved, and how they modify the cell behavior highlights the importance of this study, which could contribute to a better understanding of how Ang II may be involved in inflammatory processes of cardiovascular diseases or during vascular remodeling, even in normotensive condition.

This study hypothesizes that a non-hypertensive dose of Ang II, acting on AT1 and AT2 receptors, can trigger an early inflammatory process in small arteries, such as cardiac arteries. This process may participate in the mechanisms involved in the vascular injury or even increase any already existing damage. To test this hypothesis, the aims of this study were (1) to evaluate the expression of inflammatory markers of cardiac arteries in response to Ang II and (2) to determine by which receptor (AT1 or AT2) Ang II leads to the expression of these inflammatory markers, independently of blood pressure changes.

## Material and Methods

### Animals

The present study used adult male C57BL/6 mice provided from the animal care unit of the Department of Anatomy of the Institute of Biomedical Sciences at the University of Sao Paulo. The mice were kept in a temperature-controlled room (22°C) with a 12-h dark-light cycle and received standard laboratory chow and water ad libitum. This study was performed following the Guidelines for Ethical Care of Experimental Animals from the International Animal Care and Use Committee (National Research Council (US) Committee for the Update of the Guide for the Care and Use of Laboratory Animals, 2011). Also, it was approved and conducted according to the ethical principles established by the Ethics Committee on the Use of Animals by the Department of Anatomy of the Institute of Biomedical Sciences at the University of Sao Paulo.

To meet the study goals, three experiments were proposed: 1) to confirm that Ang II dose does not induce BP changes, 2) to evaluate a time-frame for the inflammatory response to Ang II, and 3) to study the participation of Ang II receptors in this inflammatory response.

### Confirmation of non-hypertensive dose of Ang II

Treatment with angiotensin II (Sigma®) (30ng/kg, Ang II group) and saline solution (NaCl 0.9%, control group) was performed by intraperitoneal injection with a volume of 300µL for both groups (Lima et al., 2019). Saline solution (NaCl 0.9%) was used as a vehicle for Ang II injection. Direct blood pressure (BP) and heart rate (HR) were evaluated in different experimental times after Ang II or saline injections. For carotid catheter implantation (PE-10, with an internal diameter of 0.1mm, connected to a PE-50 catheter, with an internal diameter of 0.5mm), the animals were anesthetized with isoflurane. The thinner catheter (PE-10) was introduced in the carotid artery; the thicker catheter (PE-50) was passed subcutaneously, being externalized on the back, in the cervical region. The catheters were fixed with cotton thread. Following the surgery, the animals were monitored until full recovery, and received analgesic (tramadol 40 mg/kg, every 12 hours for 2 days) and antibiotic (Benzathine Penicillin, 50 U/kg, single dose) by intramuscular injections. Forty-eight hours after complete recovery, the carotid catheter was connected to a transducer (Kent Instruments, USA) and preamplifier (Hewlet-Packard 8805C, Puerto Rico, USA). Pulse pressure was recorded on a computer by using an acquisition system (Windak 4KHz, DATAQ Instruments, Akron, OH, USA), allowing beat-to-beat analysis with 8 kHz of sampling rate per channel. Were obtained values of diastolic BP (DBP), systolic BP (SBP), mean BP (MBP), and HR for each animal.

After connecting the carotid catheter to the monitoring system, the animals were allowed to adapt to the new environmental conditions (at least 20 minutes, observing when the animals were visually calm). Basal BP and HR were measured for 20 minutes (before the injections, time 0), and after injection of saline solution (n=5) or Ang II (n=5) at the following times: 10 minutes, 30 minutes, 60 minutes, 2 hours and 6 hours.

### Time-frame for the inflammatory response to Ang II

To avoid inflammatory response inherent to surgical procedures, this experiment evaluated mice submitted only to Saline or Ang II injections (non submitted to surgeries). Heart and blood samples were collected 30 and 60 minutes and 2, 6, 12, 24, and 48 hours after Ang II injection. The control groups were evaluated after 30 minutes and 48 hours. Were analyzed 5 mice for each treatment and experimental time for histological (immunohistochemistry) and biochemical (western bloting and ELISA) analysis. *Ex vivo* analysis involved the study of local inflammatory markers by immunohistochemistry and western bloting in cardiac arteries and systemic inflammatory markers in serum by ELISA.

### Evaluation of AT1 and AT2 receptors in the inflammatory response to angiotensin II

Based on the inflammatory response to Ang II experiment, the times of 30 minutes and 12 hours were identified as moments of expression of local inflammatory markers; such times were used to study the effect of Ang II receptor blockers. To study the participation of AT1 ou AT2 receptors, six groups (n=5 for each group and time) were formed as follows: 1-control (saline solution), 2-Ang II, 3-Losartan (AT1R blocker), 4-Los+Ang II, 5-PD123.319 (AT2R blocker, ToCris®) and 6-PD+Ang II.

Ang II receptor blockers were given 30 minutes before Ang II or saline injections to allow receptor blockade before Ang II treatment. All treatments were also administered by intraperitoneal injection. The AT1 receptor blockade was done by using losartan (20mg/kg) and the AT2 receptor blockade was done by using PD123319 (15mg/kg), even in doses that do not alter blood pressure (Chen et al., 2005). For this evaluation, Ang II was also administered at a dose of 30ng/kg.

At the end of experimental times, the hearts were used for the analysis of local inflammatory markers by immunohistochemistry.

### Tissue collection and histological preparation for immunohistochemistry

At the end of each experimental time, the animals were euthanized by using an intraperitoneal overdose of ketamine (180mg/kg) supplemented with xylazine (20mg/kg). The vascular system was subsequently perfused with saline solution (0.9% NaCl) at constant pressure (80–90 mmHg) through the left ventricle followed by a buffered 4% formalin solution.

Hearts were collected and maintained in 4% buffered formalin for 24 to 48 hours allowing the complete fixation. After this period, the tissues were processed and embedded in paraplast for histological evaluation. Immunohistochemistry (IHC) was performed in four-micrometer tissue section to assess local inflammatory markers: interleukin IL-1β (abcam®9722) and IL-6 (abcam®6672), tumor necrosis factor-alpha TNF-α (abcam®6671), transforming growth factor-beta TGF-β (abcam®64715) and intercellular adhesion molecule 1-ICAM1 (abcam®25375). Briefly, ABC (streptavidin-biotin-peroxidase) method was performed using antigen retrieval by citrate buffer pH 6.0 for TGF-β, TNF-α, and ICAM-1. Slices were deparaffinized and rehydrated, following the blockade of endogenous peroxidase activity with 3% H_2_O_2_ solution for 30 minutes. After that, tissues were rinsed with phosphate-buffered saline (PBS). The primary antibodies were diluted in blocking solution (Bovine Serum Albumin 3% in PBS) and incubated in the sections for 16 hours at 4°C. The slides were washed and the biotinylated secondary antibody (Zymed Laboratories, South San Francisco, CA) was incubated for one hour at room temperature. Subsequently, the streptavidin-peroxidase complex (1: 500) was incubated for 60 minutes at room temperature. Then, 3,3’-diaminobenzidine solution (DAB, Vector Labs.) was used as a chromogen. The counterstaining was performed with hematoxylin and the slides were mounted. The negative control reactions were obtained by omitting the primary antibody in one tissue section on each slide assessed. The analyses were done blinded to the identity of experimental groups. The evaluation of immunohistochemistry was performed as previously described (Lacchini et al., 2009; Lima et al., 2015). Briefly, the semiquantitative analysis used a score of 0 to 4, where 0 represents no staining, 1=weak staining, 2= weak to moderate staining, 3= moderate staining, and 4=intense staining. Three to five sections per heart were evaluated, examining at least five cardiac arteries per animal (each heart was represented by the mean from all analyzed arteries).

### Tissue and blood collection to study inflammatory markers by Western Blot and ELISA

At the end of experimental times, mice were euthanized as described previously. All time-frame groups (n=5 for treatment and time) were assessed to determine whether Ang II was able to change cardiac protein expression (western blot) and if it was also capable of inducing systemic effects on inflammatory markers (ELISA). For these evaluations, blood samples were collected directly from the chest cavity after the right atrium had been cut. Following, the heart was collected and immediately frozen in liquid nitrogen for further western blot analysis.

### Protein quantification in the heart

Western blot analysis was performed by lysing hearts in RIPA buffer (1mMEDTA, 1mMEGTA, 2mMMgCl2, 5mMKCl, 25mM Hepes, pH 7.5, 2mM DTT, 1mM PMSF, 0.1% Triton X-100, and 1:100 cocktail of protease inhibitors) and stirring for 30 minutes at 4∘C. Homogenates were centrifuged (10000×g for 10min, 4∘C), and the supernatant collected. Tissue lysate (50μg) was heated up in a sample buffer (200mM Tris-HCl, pH 6.8, 40% glycerol, 8% sodium dodecyl sulphate, 0.1% DDT, and 0.4% bromophenol blue) at 100∘C for 5 minutes. The samples were run in a sodium dodecyl sulfate-polyacrylamide gel electrophoresis (SDS-PAGE), transferred to a nitrocellulose membrane *Hybond-C Extra* (GE Healthcare) in transfer medium (0.025MTris, 0.192Mglycine, 0.1% SDS, and 20% methanol). Membranes were washed (3 times in 1X TBS-T, 5 minutes), blocked (5% BSA, 3 hours), washed in TBS-T (3 times), and incubated overnight with antibody anti-TGF-*β* (1:500, sc-146, Santa Cruz Biotechnology) or anti-IL-6 (1:500, ab6672, Abcam). Membranes were exposed in *ECL WB Detection Reagents* (GE Healthcare) and revealed in *Image Quant LAS 4000 mini* (GE Healthcare) equipment. Protein bands were quantified by optical densitometry using *ImageJ* software (version 1.32j, NIH). GAPDH expression (1:1000, Santa Cruz) was used to normalize the results.

### Protein quantification in the serum

Systemic evaluation of inflammatory markers (IL-6 and TNFα) was made by enzyme-linked immunosorbent assay (ELISA) in serum. ELISA evaluation used Mouse TNF-alpha/TNFSF1A DuoSet and Mouse IL-6 DuoSet according to the manufacturer’s specifications. Results were obtained by using the ELISA reader Spectramax PLUS v3 ROM with a wavelength of 450nm. The reader is coupled to a microcomputer and data were analyzed by software SoftMax Pro 5.2 for the calculations.

### Statistical Analysis

Values are expressed as mean±standard deviation. The hemodynamic and western blot data were statistically evaluated by one-way variance analysis (ANOVA) complemented using Bonferroni’s test. As immunohistochemistry evaluation for time curve and AT1R blockers was made by using score analysis, the results were treated by the Kruskal-Wallis test complemented by the test of Dunns. For all analyses, the significance level of p≤0.05 was adopted.

## Results

### Confirming of a non-hypertensive dose of Ang II

Direct blood pressure assessment was performed to confirm the absence of hypertensive effect mediated by injected dose of Ang II. Thus, variations in inflammatory markers would not be related to blood pressure changes, but rather to a direct effect of Ang II. Table 1 shows diastolic blood pressure (DBP), systolic blood pressure (SBP), mean blood pressure (MBP) and heart rate (HR). As it is demonstrated, the treatments did not affect blood pressure or heart rate even after 10 or 30 minutes until 6 hours after injection, corroborating with previous studies (Lacchini et al., 2009; Lima et al., 2019).

**Table 1.** Blood pressure (BP) and heart rate measurements: diastolic BP (DBP), mean BP (MBP), systolic BP (SBP) and heart rate (HR) were measured in Saline and Ang II treated mice. The same animal was recorded at basal time (baseline) and 10, 30, 60 120 minutes and 6 hours after injections.

|  | Saline<br>(n=5) |  |  |  | Ang II<br>(n=5) |  |  |  |
| --- | --- | --- | --- | --- | --- | --- | --- | --- |
|  | DBP<br>(mmHg) | SBP<br>(mmHg) | MBP<br>(mmHg) | HR<br>(bpm) | DBP<br>(mmHg) | SBP<br>(mmHg) | MBP<br>(mmHg) | HR<br>(bpm) |
| Baseline | 103,7±4,8 | 135,2±1,2 | 120,3±3,4 | 563±144 | 89,9±8,4 | 127,4±4,8 | 108,8±6,7 | 528±57 |
| 10 min | 105,8±0,7 | 135,1±2,3 | 121,5±0,8 | 626±186 | 91,8±10,2 | 131,4±13,4 | 111,9±11,4 | 538±59 |
| 30 min | 100,3±1,2 | 128,6±3,0 | 115,8±0,6 | 587±108 | 86,9±6,2 | 120,9±4,7 | 104,6±5,4 | 545±56 |
| 60 min | 100,2±4,2 | 126,1±3,1 | 114,7±3,5 | 655±81 | 84,8±4,7 | 118,7±6,7 | 102,4±3,9 | 563±87 |
| 120 min | 90,7±2,0 | 122,3±8,8 | 107,8±3,1 | 583±123 | 86,9±14,3 | 122,2±11,9 | 104,9±12,8 | 552±65 |
| 6 hours | 79,9±2,7 | 115,7±7,1 | 99,2±3,7 | 537±153 | 80,9±11,6 | 114,4±7,2 | 98,4±9,5 | 510±61 |

Statistical analysis did not show differences between treatments for all parameters. n=5 per treatment and time

### Time-frame of the inflammatory response to Ang II

#### Analysis of local inflammatory markers

The local inflammatory markers evaluated were: TGF-β, TNF-α, IL-1β, IL-6 and ICAM-1. This analysis in sequential times after treatment allowed us to have a temporal idea of the phenomenon and to understand how the artery responds to the stimulus. Briefly, it was possible to identify two types of inflammatory response: one earlier (30-120 minutes) and one later (6 to 48 hours).

Figure 1 shows the analysis by score obtained in inflammatory markers with an earlier response. It is possible to observe a fast response of the cardiac arteries, showing an increase in TGF-β staining from 30 to 120 minutes after the injection with Ang II (Figure 1A). After 120 minutes, the positive staining resembles that seen in the Saline groups. A similar response was observed when evaluating the positive label for IL-6 (Figure 1B), showing an increase in this protein from 20 to 60 minutes. Both TGF-β and IL-6 showed early increases, which can be attributed to a direct action of Ang II.

**Figure 1.**
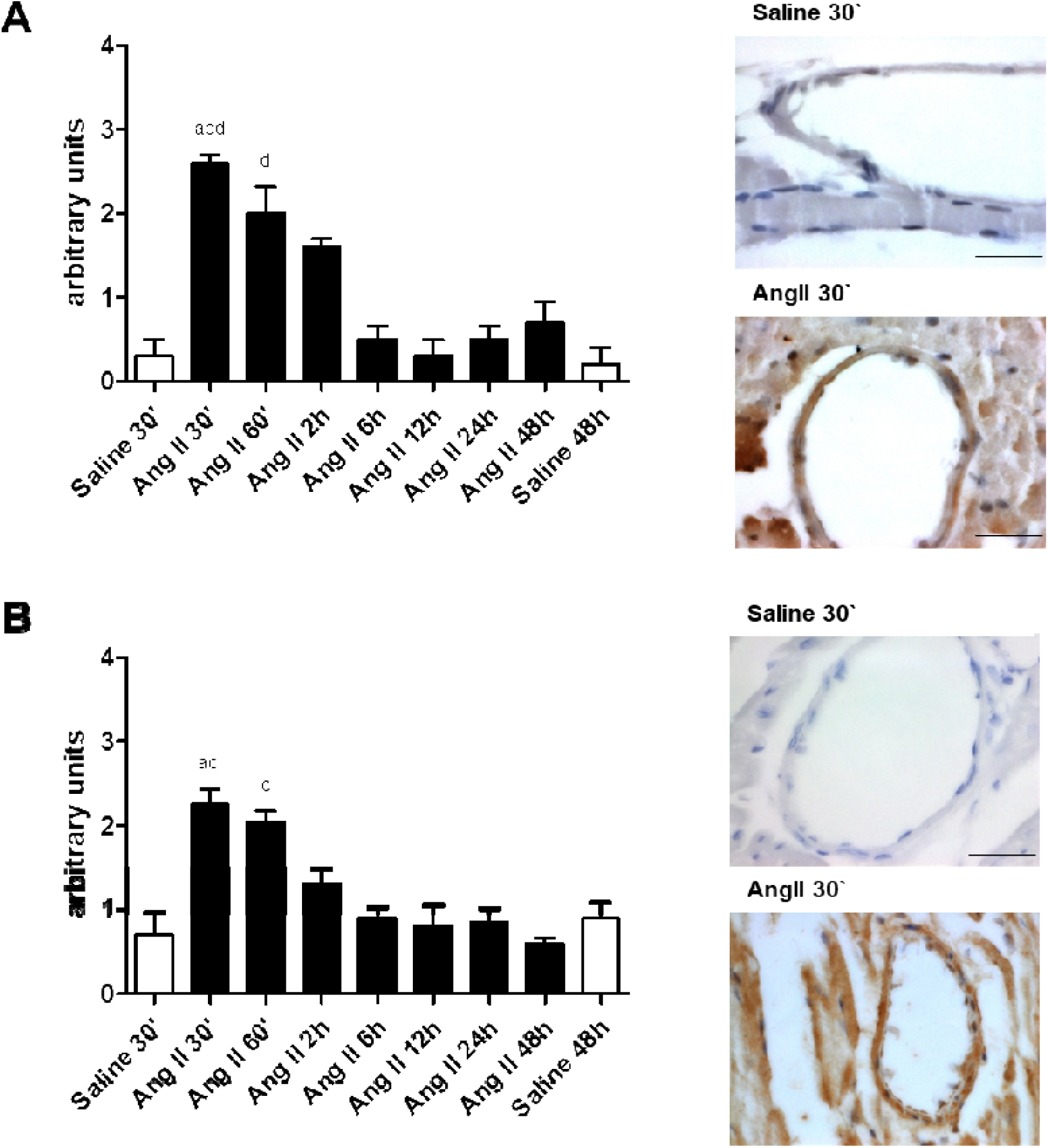
Time curve in response to Saline or Ang II injection, showing score staining for TGF-β (A) and IL-6 (B) on left; on right side are presented representative images comparing basal condition (Saline) and higher response (30’ after Ang II injection). *p≤0.05 compared to Saline 30’ (a), Ang II 12h (b), Ang II 48h (c) and Saline 48h (d). *Scale bar=0,4 µm*.

On the other hand, IL-1β and ICAM-1 show later increase. IL-1β increased in cardiac arteries from 2 to 24 hours after the injection of Ang II (Figure 2A), ICAM-1 was more produced only 6 to 12 hours after the stimulus (Figure 2B). TNF-α did not show any significant changes at all studied times (Figure 2C).

**Figure 2.**
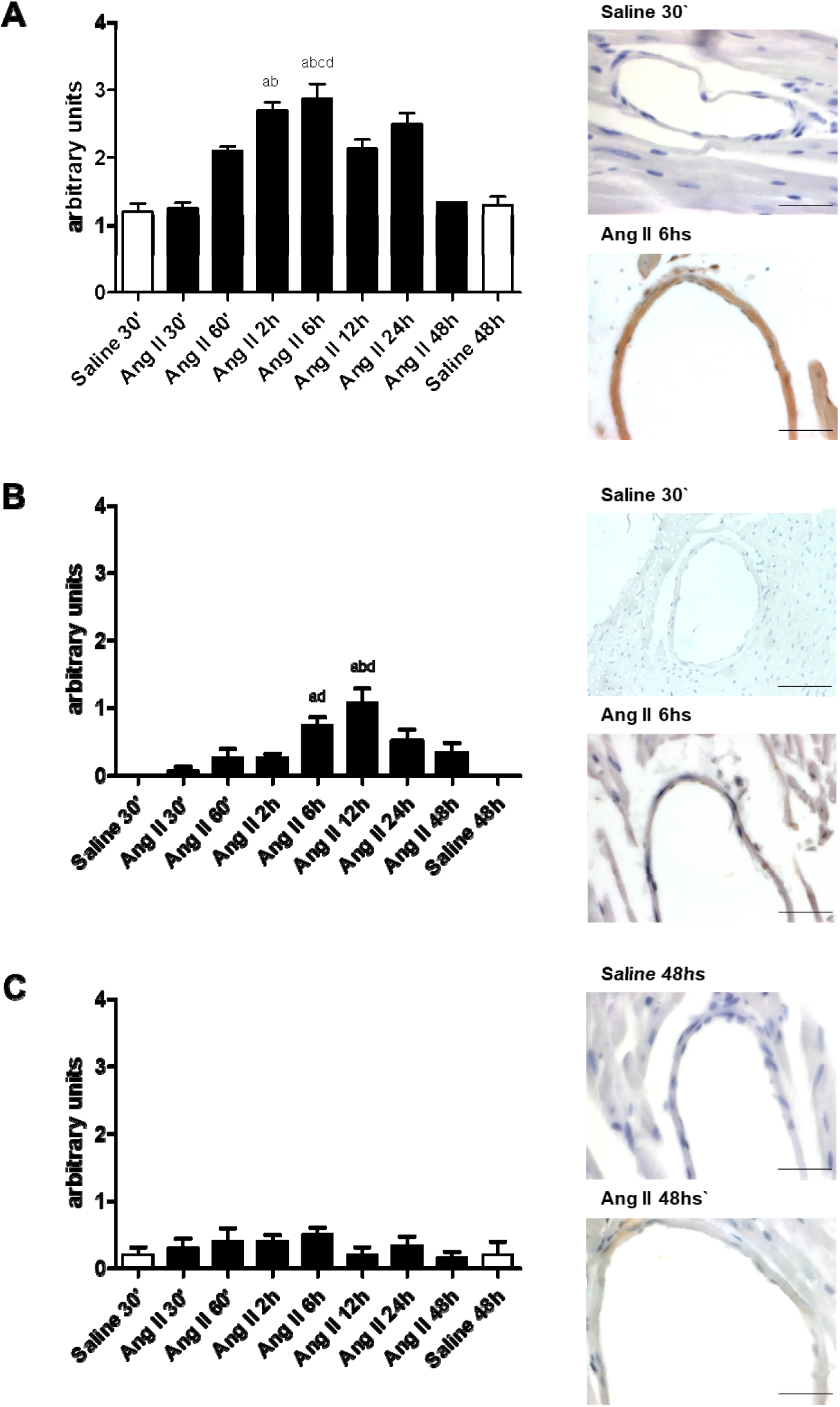
Time curve in response to Saline or Ang II injection, showing score staining for IL-1β (A), ICAM-1 (B) and TNF-α (C) on left; on right side are presented representative images comparing basal condition (Saline) and higher response (6 or 48 hours after Ang II injection). *p≤0.05 compared to Saline 30’ (a), Ang II 30’ (b), Ang II 48h (c) and Saline 48h (d). *Scale bar=0,4 µm*.

#### Analysis of cardiac inflammatory markers

As the inflammatory markers that showed an early change in the cardiac arteries were TGF-β and IL-6, it was considered important to verify whether this response would be only vascular or would involve the entire cardiac tissue. Therefore, the protein expression of such markers was measured by Western blot. Interestingly, an early cardiac response to Ang II was found for both markers. Figure 3 shows the percentual (/GAPDH ratio) increase in protein expression of TGF-β after 120 minutes and IL-6 after 60 minutes. This result suggests that the inflammatory response mediated by a small dose of Ang II may induce a fast tissue response, including vessels and the entire tissue.

**Figure 3.**
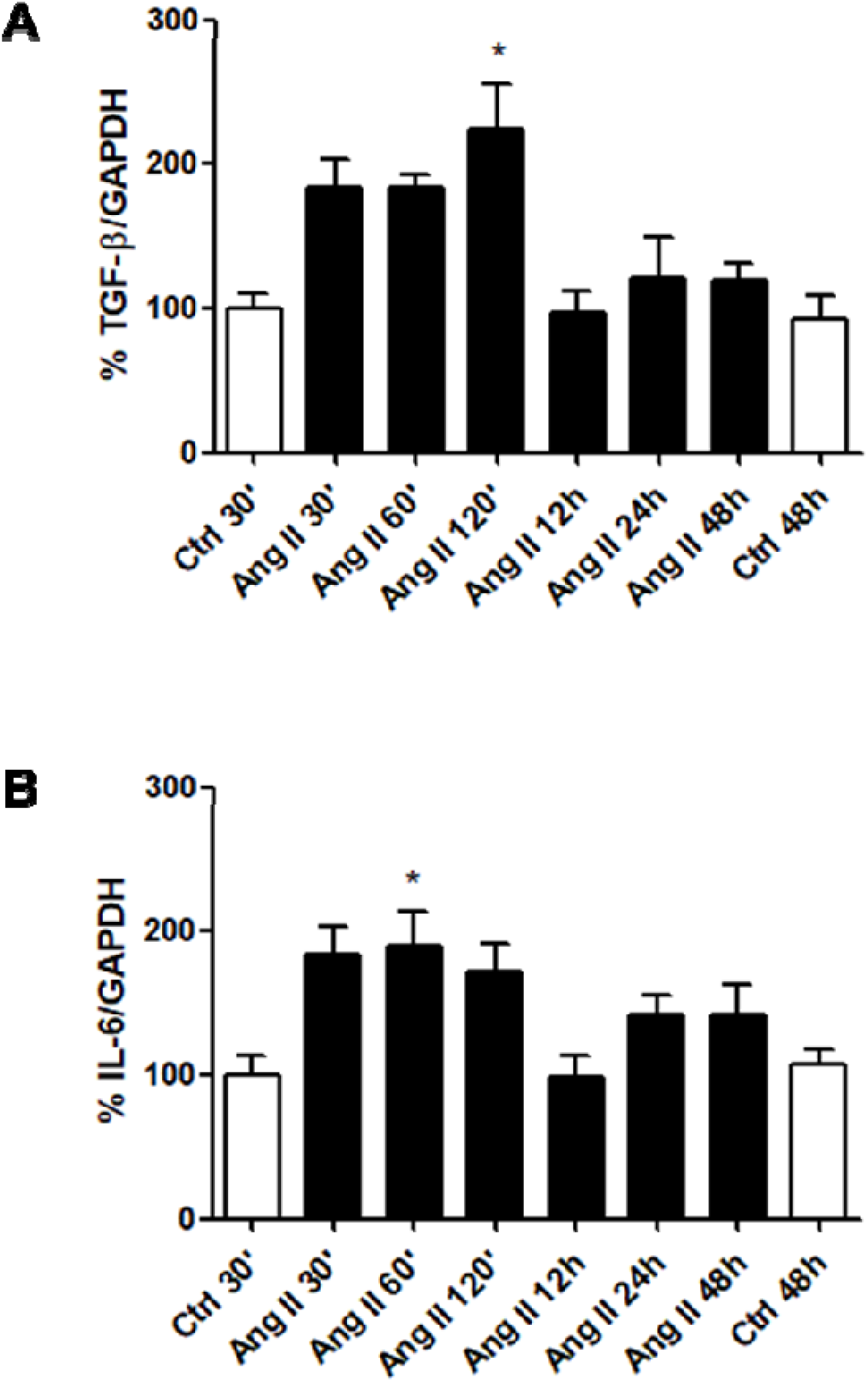
Cardiac expression of TGF-β and IL-6. The results are presented as percentage of protein/GAPDH ration in all times after Saline or Ang II injection. *p≤0.05 compared to Saline and other Ang II groups.

#### Evaluation of systemic inflammatory markers by ELISA

Systemic inflammatory markers were quantified in serum by ELISA. Table 2 shows the time curve obtained for TNF-α and IL-6. Although it was possible to observe an increase in the serum concentration of TNF-α, this was not statistically significant, corroborating the local inflammatory analysis. The concentration of IL-6, on the other hand, increased significantly 60 minutes after the injection of Ang II, coinciding with the results observed in cardiac arteries and heart, as previously described.

**Table 2.** Systemic (serum) TNF-α and IL-6 measured by ELISA. Each group was designed for a treatment (saline or Ang II) and the blood was collected after the time specified.

| Treatment | TNF-α (pg/mL) | IL-6 (pg/mL) |
| --- | --- | --- |
| Saline 30' | 0 ± 0 | 0 ± 0 |
| Ang II 30' | 3,27 ± 7,32 | 1,60 ± 3,59 |
| Ang II 60' | 6,28 ± 14,04 | 29,21 ± 11,86 * |
| Ang II 120' | 0 ± 0 | 1,65 ± 3,70 |
| Ang II 6hs | 0 ± 0 | 0 ± 0 |
| Ang II 12hs | 0 ± 0 | 0 ± 0 |
| Ang II 24hs | 0 ± 0 | 0,56 ± 1,26 |
| Ang II 48hs | 0 ± 0 | 1,17 ± 2,63 |
| Saline 48hs | 0 ± 0 | 0 ± 0 |

\*p ≤ 0.05 compared to Saline groups and other times of Ang II; n=5 per treatment and time

### Role of AT1 and AT2 receptors in the local inflammatory response to angiotensin II

The analysis of the possible role of Ang II receptors on the local expression of inflammatory markers was based on the time curve, where we verified that TGF-β and IL-6 present early changes (the time of 30 minutes after Ang II injection was chosen), while IL-1β and ICAM-1 show later changes (the time of 12 hours after injection was chosen).

The early analysis (30 minutes) after treatment reinforced the previous result that Ang II increases TGF-β and IL-6 expression in cardiac arteries. As can be seen in Figure 4, Losartan was able to inhibit the increase of TGF-β (Figure 4A) and IL-6 (Figure 4B), suggesting that AT1 receptor is mediating such an increase of both cytokines. On the other hand, the concomitant treatment of Ang II with the AT2R blocker PD123.319 shows the same inhibition of the TGF-β expression in cardiac arteries, indicating a possible role of both AT receptors in this response.

**Figure 4.**
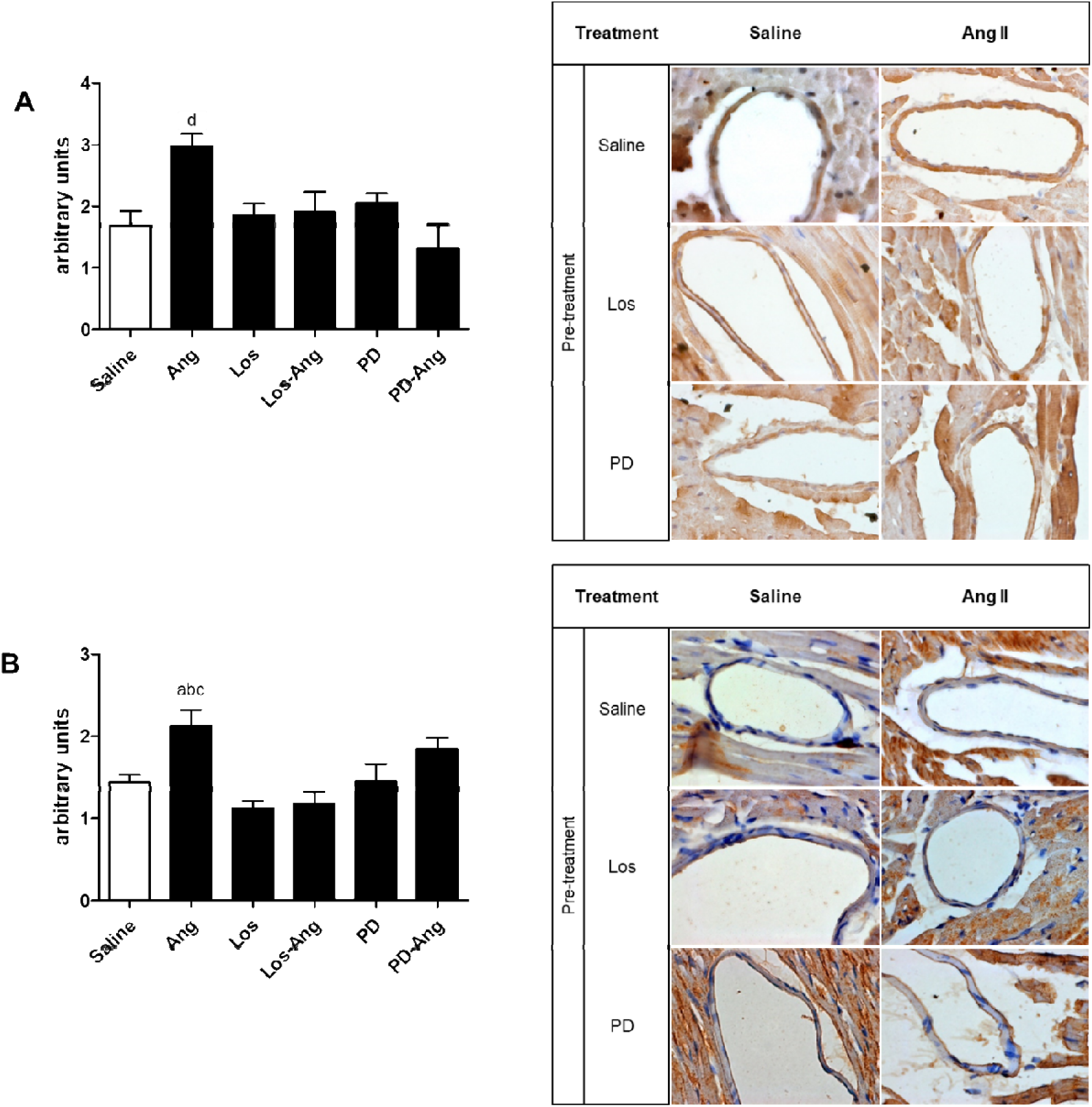
Study of the participation of AT receptors on local expression of TGF-β (A) and IL-6 (B) in early inflammatory response (30 minutes after treatments). On left is presented the score of staining in each group, and on right is presented a panel with representative images of the results obtained after treatments and ATR blockings. *p≤0.05 compared to Saline (a), losartan (b), los+Ang (c) and PD+Ang (d). Magnification: 1000x

The late evaluation supports the previous results, showing that Ang II increases IL-1β and ICAM-1 immunostaining in cardiac vessels (Figure 5). On the other hand, we observed inhibition in the expression of these cytokines in the losartan-combined treatment suggesting that Ang II effect is also related to the AT1 receptor. PD123.319 was not efficient in reducing IL-1β, which indicates that the AT2 receptor possibly does not participate in the modulation of this cytokine performed by Ang II. Interestingly, PD123.319 alone or combined with Ang II increased ICAM-1 levels in the cardiac arteries. Although these results are not statistically significant, it is debatable a possible action of AT2 antagonizing the effects of AT1 in these vessels.

**Figure 5.**
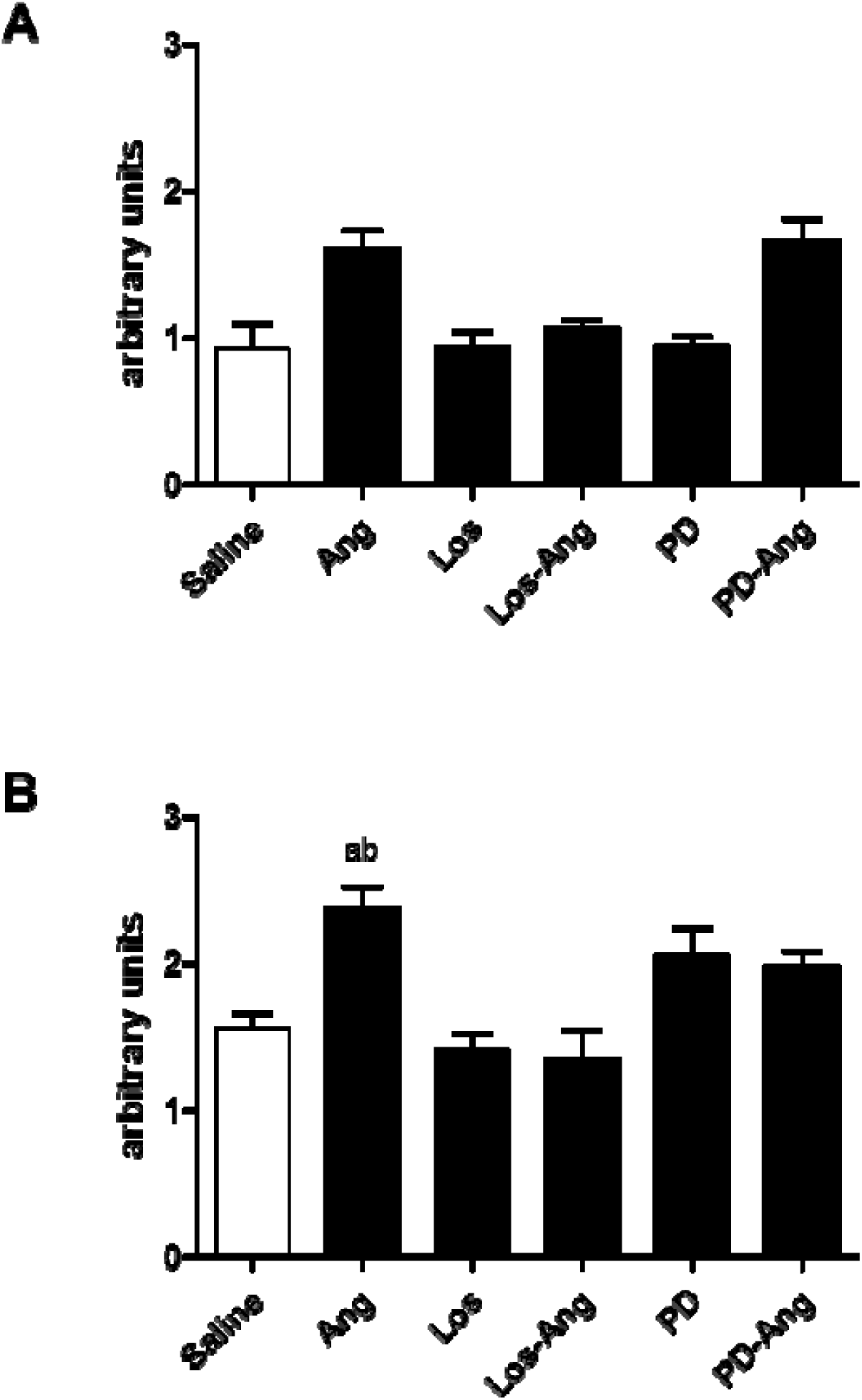
Study of the participation of AT receptors on local expression of IL-1β (A) and ICAM-1 (B) in late inflammatory response (12 hours after treatments). *p≤0.05 compared to losartan (a) and los+Ang (b). Magnification: 1000x

## Discussion

This study aimed to determine if Ang II is capable of inducing an inflammatory response in cardiac arteries even under normal values of blood pressure. It is well known that BP elevation *per se* can stimulate tissues to produce inflammatory cytokines as a repair mechanism (Guzik & Touyz, 2017). The concept of angiotensin II with non-hemodynamic functions arose from studies conducted mainly in the late 1990s (Das, 2005a, 2005b). Specifically, the proinflammatory effect of Ang II started correlating the action of Ang II with the production of reactive oxygen species leading to the production of inflammatory cytokines, and the expression of adhesion molecules (Heinrich et al., 2003; Sadoshima, 2000; Schieffer et al., 2000; Ushio-Fukai et al., 1999). This idea was supported by the demonstration that AT1 receptor blockers can reduce inflammation, leading to atherosclerotic plaque stabilization (Cipollone et al., 2004).

Besides the proinflammatory action, it is important to consider that studies evaluating the effect of angiotensin II are usually made in animal models of hypertension. Thus, this hypertensive state can lead to a cardiovascular inflammatory condition, creating a confounding complementary factor. For example, hypertensive patients have an increase in adhesion molecules in serum and inflammatory cells (Ruiz-Ortega et al., 2001, 2007) but this fact can be directly associated with the inflammatory actions of Ang II. Therefore, this study addressed the inflammatory effect of angiotensin II, seeking to isolate the hemodynamic aspect using a dose that is not capable of altering BP. Trying to respond to this important question, the present study intended to isolate the Ang II stimulus from BP variations. Here, we demonstrate that a low dose of Ang II (30ng/kg) is unable to induce BP changes and leads to an early local and systemic inflammatory response.

Regarding the effect of Ang II, we believe that it presents a fast response and the entire result comes from the action of Ang II itself. We presume that Ang II, even being metabolized very quickly, is responsible for creating a chain-reaction mediating an inflammatory response. Apart from this, if we consider the fast metabolization of Ang II, which could not be time effective in promoting a systemic response, even the chronic treatment with osmotic mini-pumps, would also not be effective once to reach the circulatory system, the Ang II needs to be absorbed by the capillaries formed around the osmotic pumps, thus the effect of the Ang II would be reduced by the use of the osmotic mini-pumps. However, it has been shown that the administration of a low or high dose of Ang II by a osmotic mini-pumps is very effective (Daugherty et al., 2000; Guan et al., 2017; Guzik & Touyz, 2017). For this reason, we believe that the intraperitoneal administration of a single dose of Ang II reaches the circulatory system when in contact with a well-developed venous network that absorbs this Ang II very quickly, transporting it via the hepatic portal system to the heart. Therefore, a cardiac effect is expected, as was observed in this study. To better understanding whether the injection itself would have a stress-based effect, we also administered a saline solution to a control group. It should be noted that for ethical reasons, and also to reduce the number of animals, we did not perform the entire time-frame for the control group. We decided to analyze this effect in the times 30 minutes and 48 hours. Furthermore, as we did not observe a significant difference in the levels of IL6 and TNF-α by immunostaining, we did not consider relevant analyzing these inflammatory markers in the saline group by ELISA.

In this study, we observed no significant changes in blood pressure and heart rate comparing periods of 10 minutes to 6 hours after Ang II injection with a basal period, which agree with a previous study (Lima et al., 2019). This study also confirmed the non-hypotensive dose of Losartan (AT1 receptor blocker) and the combination of Losartan and Ang II, as previously demonstrated (Lacchini et al., 2009). The fact that no significant change was observed in blood pressure or heart rate suggests that the Ang II-induced responses are independent of hemodynamic alterations.

### Study of the time-frame in response to angiotensin II

The time curve allowed us to identify moments when the inflammatory response is activated, both locally and systemically. This effect occurred both locally (cardiac arteries and cardiac tissue) and systemically (serum), reflecting the importance of a small Ang II increase in cardiovascular regulation. The evaluation of inflammatory markers allowed to identify early and late inflammatory responses to Ang II.

TGF-β and IL-6 markers already had an increase in vascular and tissue expression from 30 to 120 minutes after the Ang II stimulus. Moreover, we verified a significant increase of IL-6 in serum 60 minutes after the injection of Ang II. Some studies show the fast expression of IL-6 in response to different types of stress such as infection or exercise. In chronic inflammation, the expression of TNF-α precedes IL-6; however, the stimulus without inflammation expresses IL-6 without being preceded by TNF-α (Pedersen & Febbraio, 2008), corroborating with the results obtained in the present study.

IL-6 is produced by a large number of cell types, and its release includes the presence of other pro-inflammatory cytokines, oxidation of lipoproteins, and the activation of extracellular matrix metalloproteinases (Schuett et al., 2009). It is known that Angiotensin II is capable of inducing the vascular production of pro-inflammatory cytokines such as IL-6 and TNF-α (Schieffer et al., 2000). In this case, Angiotensin II would lead to an increase in the production of reactive oxygen species, inducing an increase in IL-6 production (Wassmann et al., 2004), and creating a chain-reaction in response to Angiotensin II, promoting an increase in IL -6, oxidative stress and vascular injury.

It has been shown that TGF-β is a multifunctional protein, participating in the regulation of cell division, differentiation, migration, cell adhesion, extracellular matrix production, and is involved in several diseases including cardiovascular disease (Ruiz-Ortega et al., 2007). It should be noted that TGF-β protein is synthesized as inactive but has within its structure the latency-associated peptide (LAP). This protein interacts with protein binding of latent TGF-β (LTBP) which is anchored to the extracellular matrix. This TGF-β is activated by proteolytic cleavage by plasmin, microenvironment acid, and matrix metalloproteinases (MMP-2 and 9) (Annes et al., 2003). In vitro studies show that Ang II stimulates the gene expression and activation of TGF-β in vascular smooth muscle cells (Weigert et al., 2002). However, the fast response observed in TGF-β evaluation suggests that it is probably anchored to proteins in the extracellular matrix and is released under-stimulation. In this study, Ang II was able to promote a rapid release of TGF-β in cardiac arteries. This idea corroborates a previous study, showing a similar pattern of TGF-β release in the aorta of normotensive mice (Lima et al., 2019). Moreover, by participating in the balance between inflammation and extracellular matrix deposition, the local imbalance in TGF-β can lead to the development of early vascular lesions and atherosclerosis (Ruiz-Ortega et al., 2007).

The results for both IL-1β, TNF-α, and ICAM-1 are consistent with the observation of an early increase in IL-6 both locally and in serum. In response to a harmful stimulus, the body triggers an innate nonspecific reaction activating an acute protein reaction (APR), as IL-1β, IL-6, and TNF-α, involved in local inflammation (Schuett et al., 2009). IL-6 is the main mediator of such mechanism [Heinrich et al., 1998] acting on the expression of other cytokines and inflammatory mediators. It has been demonstrated that IL-6 has an inhibitory effect on the production of TNF-α (Mizuhara et al., 1994) which corroborates the results found in this study because, unlike a situation of infection, the injection of Ang II led to increased IL-6 without being preceded by an increase in TNF-α. In contrast, IL-6 stimulates the release of IL-1β maintaining the inflammatory response as observed. This response was maintained between 2 and 24 hours and possibly declined at 48 hours due to a lack of stimulus (in this case, the injection of Ang II). The maintenance of an inflammatory stimulus leads to the facilitation of adhesion and cell migration in the target tissue. In this context, the observation of increased staining of ICAM-1 suggests also a proinflammatory action, and this is corroborated by the fact that both IL-6 and IL-1β are stimulating factors as ICAM-1 (Zhang et al., 2009).

### Study of the effect of Ang II receptors on inflammatory response

Although this study showed that Ang II, independent of hemodynamic changes, can trigger a local inflammatory response in cardiac arteries, it is questionable whether this action is mediated by the direct action of its receptors or if it can be an indirect mechanism that acts on this response. Thus, the study of the receptor blockade of Ang II allowed us to observe the possible participation of each AT receptor on the inflammatory process triggered by Ang II.

As can seeing in results, the injection of Ang II leads to a fast increase in TGF-β and IL-6 in cardiac arteries. Also, it can increase IL-1β and ICAM-1 after 12 hours in such vessels. Is has been well described that this Ang II effect is related to its action on AT receptors (Ferrario & Strawn, 2006; Sadoshima, 2000; Sakuta et al., 2010), but is always associated with different pathologies. The present study shows that this effect is either non-associated to pathologies or hemodynamic changes.

The proinflammatory effect of Ang II-mediated by the AT1 receptor is confirmed by blockade with Losartan. As Losartan abolished the effect of Ang II, we may consider that AT1 acts directly to increase vascular TGF-β and IL-6 after 30 minutes. However, the experimental design of this study does not allow us to assert that the effects on IL-1β and ICAM-1 12 hours after Ang II injection are directly determined by AT1 activation. This expression may be given indirectly after the activation of AT1. In this context, it opens a new possibility for study, analyzing the mechanisms by which activation of AT1 leads to increased expression of late inflammatory mediators, such as IL-1β and ICAM-1. The effects of the AT2 receptor are recognized and widely studied as AT1 antagonists. Thus, AT2 has classically exerted antiproliferative and antiinflammatory actions, among others (Ferrario & Strawn, 2006; Horiuchi et al., 1999). Furthermore, we found that AT2 does not always participate in Ang II-related effects, restricting the actions of Ang II to those mediated by AT1. In this sense, we found that the AT2 receptor seems to have an AT1 antagonism action, especially related to IL-1β.

### Study limitations

In this study, the observed effects of Ang II on inflammatory markers in cardiac arteries were associated with AT1 receptors. Although the protein expression has been analyzed, the results obtained represent a set mostly composed of cardiomyocytes, diluting the specific analysis of the cardiac arteries. However, this evaluation allowed us to verify that the cardiac tissue itself responded to the treatment, producing an inflammatory response.

Another limitation that we consider for this work is the difficulty of working with small groups of animals, since many experimental groups are evaluated, and the reduction strategies of the ethics committee provide for the use of a minimum number of animals. Studies using immunohistochemistry can show a lot of variation within an experiment and attention should be maximum, especially with small groups.

Furthermore, the results related to the AT2 receptor can be confused with results from the other groups. The idea of the interaction between the two Ang II receptors has existed for over 10 years but the topic is controversial. It is believed that the clinical effects of AT1 receptor antagonists may be due to the blockade of AT1 and AT2 stimulation whereas Ang II would act on the AT2 receptor preferentially when AT1 is blocked (Horiuchi et al., 1999). The real contribution of the AT2 receptor antagonizing the effects of AT1 can be better understood in studies with dual blockade of the receptors. This opens new perspectives in understanding the role of both AT1 and AT2 in the physiological responses to Ang II.

## Conclusion

It is noteworthy that the effect of Ang II on the early or late expression of inflammatory markers in the arteries of healthy individuals may not represent risks; however, we do not know whether the time and/or the repetition of these insults can participate in the development of the cardiovascular disease. Considering the importance of understanding the adaptive processes in the transition between normotensive to hypertension, the understanding of how certain changes occur in the absence of established hypertension can be fundamental to determine how and when certain changes can be reversed. In conclusion, this study showed for the first time that angiotensin II might trigger an inflammatory process mediated by the AT1 receptor on cardiac arteries, in dose small enough to keep unchanged the blood pressure. AT2 receptor seems to be involved in the counterbalance of AT1R during modulation of IL-1b production in cardiac arteries.

## Acknowledgment

Coordenação de Aperfeiçoamento de Pessoal de Nível Superior (CAPES) and Fundação de Amparo à Pesquisa do Estado de São Paulo (FAPESP, process n° 2007/57416-3 adn 2009/15693-6) for financial support.

